# Cellobiose consumption uncouples extracellular glucose sensing and glucose metabolism in *Saccharomyces cerevisiae*

**DOI:** 10.1101/076364

**Authors:** Kulika Chomvong, Daniel I. Benjamin, Daniel K. Nomura, Jamie H.D. Cate

**Affiliations:** Department of Plant and Microbial Biology, University of California, Berkeley, CA 94720; Department of Nutritional Sciences and Toxicology, University of California, Berkeley, CA 94720; Department of Molecular and Cell Biology, University of California, Berkeley, CA 94720; Department of Chemistry, University of California, Berkeley, CA 94720; Physical Biosciences Division, Lawrence Berkeley National Laboratory, Berkeley, CA 94720

**Keywords:** cellobiose, glucose sensors, metabolomics, PMA1

## Abstract

Glycolysis is central to energy metabolism in most organisms, and is highly regulated to enable optimal growth. In the yeast *Saccharomyces cerevisiae*, feedback mechanisms that control flux through glycolysis span transcriptional control to metabolite levels in the cell. Using a cellobiose consumption pathway, we decoupled glucose sensing from carbon utilization, revealing new modular layers of control that induce ATP consumption to drive rapid carbon fermentation. Alterations of the beta subunit of phosphofructokinase (*PFK2*), H^+^-plasma membrane ATPase (*PMA1*), and glucose sensors *(SNF3, RGT2)* revealed the importance of coupling extracellular glucose sensing to manage ATP levels in the cell. Controlling the upper bound of cellular ATP levels may be a general mechanism used to regulate energy levels in cells, via a regulatory network that can be uncoupled from ATP concentrations under perceived starvation conditions.

**Importance:** Living cells are fine-tuned through evolution to thrive in their native environments. Genome alterations to create organisms for specific biotechnological applications may result in unexpected and undesired phenotypes. We used a minimal synthetic biological system in the yeast *Saccharomyces cerevisiae* as a platform to reveal novel connections between carbon sensing, starvation conditions and energy homeostasis.

## Introduction

Most microorganisms favor glucose as their primary carbon source, as reflected in their genetic programs hard-wired for this preference. Central to carbon metabolism is glycolysis, which is finely tuned to the dynamic state of the cell due to the fact that glycolysis first consumes ATP before generating additional ATP equivalents. To avoid catastrophic depletion of ATP, the yeast *Saccharomyces cerevisiae* has evolved a transient ATP hydrolysis futile cycle coupled to gluconeogenesis (1). Glycolysis in yeast is also tightly coupled to glucose transport into the cell, entailing three extracellular glucose sensing mechanisms and at least one intracellular glucose signaling pathway (2).

Synthetic biology and metabolic engineering of yeast holds promise to convert this microorganism into a “cell factory” to produce a wide range of chemicals derived from renewable resources or those unattainable through traditional chemical routes. However, many applications require tapping into metabolites involved in central carbon metabolism, a daunting challenge as living cells have numerous layers of feedback regulation that fine-tune growth to changing environments. Cellular regulation evolved intricate networks to maintain and ensure cell survival. For example, *S. cerevisiae* has evolved to rapidly consume high concentrations of glucose through fermentation, while repressing the expression of other carbon consumption pathways, an effect termed glucose repression. When perturbed genetically, regulatory networks such as those in glucose repression often generate undesirable or unexpected phenotypes.

For yeast to be useful in producing large volumes of renewable chemicals or biofuels, it will be important to expand its carbon utilization to include multiple sugars in the plant cell wall. One promising approach that helps overcome glucose repression and allows simultaneous utilization of different sugars is cellobiose consumption (3). Cellobiose is a disaccharide with two units of glucose linked by a ß-1,4 glycosidic bond. Cellobiose consumption using a minimal additional pathway in yeast—a cellodextrin transporter (CDT-1) and intracellular ß-glucosidase (4)–avoids glucose repression by importing carbon in the form of cellobiose instead of glucose. The cellodextrin transporter allows cellobiose to enter the cell where it is hydrolyzed to glucose and consumed via the native glycolytic consumption pathway. By moving glucose production into the cell, the *Neurospora crassa*-derived cellobiose consumption pathway is nearly the minimal synthetic biological module imaginable in *S. cerevisiae*, comprised of just two genes. Nevertheless, in *S. cerevisiae* the cellobiose consumption pathway is inferior to consumption of extracellular glucose in terms of rate and results in a prolonged lag phase (5). Previous efforts to understand the impact of cellobiose consumption on the physiology of *S. cerevisiae* using transcriptional profiling revealed that cellobiose improperly regulates mitochondrial activity and amino acid biosynthesis, both of which are tightly coupled to the transition from respiration to fermentation (5).

Since glycolytic enzymes are regulated mostly at the post-transcriptional level (6), we probed cellobiose consumption in *S. cerevisiae* at the metabolite level. We found that key metabolites in glycolysis are highly imbalanced, leading to low flux through glycolysis and slow fermentation. We also found that excess ATP levels drive the imbalance, and identified a new potential regulatory role of glucose sensors in cellular ATP homeostasis.

## Results

### Metabolite profiling of cellobiose utilizing *S. cerevisiae*

*S. cerevisiae* cells engineered with the cellobiose consumption pathway exhibits a prolonged lag phase, with decreased growth and carbon consumption rates in comparison to when glucose is provided (Figure S1A) (5). We hypothesized that cellobiose consumption results in an ATP deficit in glycolysis relative to glucose, due to the fact that the cellodextrin transporter (CDT-1) in the cellobiose utilizing pathway is a proton symporter, requiring ATP hydrolysis for cellobiose uptake (7). Moreover, under anaerobic conditions, ATP production is limited to substrate-level phosphorylation, further restricting ATP availability. We measured the steady-state concentrations of ATP and other metabolites in central carbon metabolism in yeast fermenting cellobiose compared to glucose. Of the 48 compounds analyzed, the abundance of 25 compounds was significantly different between cellobiose and glucose-fed samples (Figure S1B). Surprisingly, ATP levels increased by 6-fold in the cellobiose grown cells (Figure 1A). The relative abundance of compounds in glycolysis—fructose 6-phosphate (F6P), glucose 6-phosphate (G6P), glucose and pyruvate—increased by 444-, 81-, 7- and 3-fold, respectively, while that of phosphoenolpyruvate (PEP) decreased by 2-fold (Figure 1A, B). These results suggest that the yeast cells underwent drastic physiological changes, reflected in metabolite levels, when cellobiose was provided in place of glucose.

**Figure 1.**
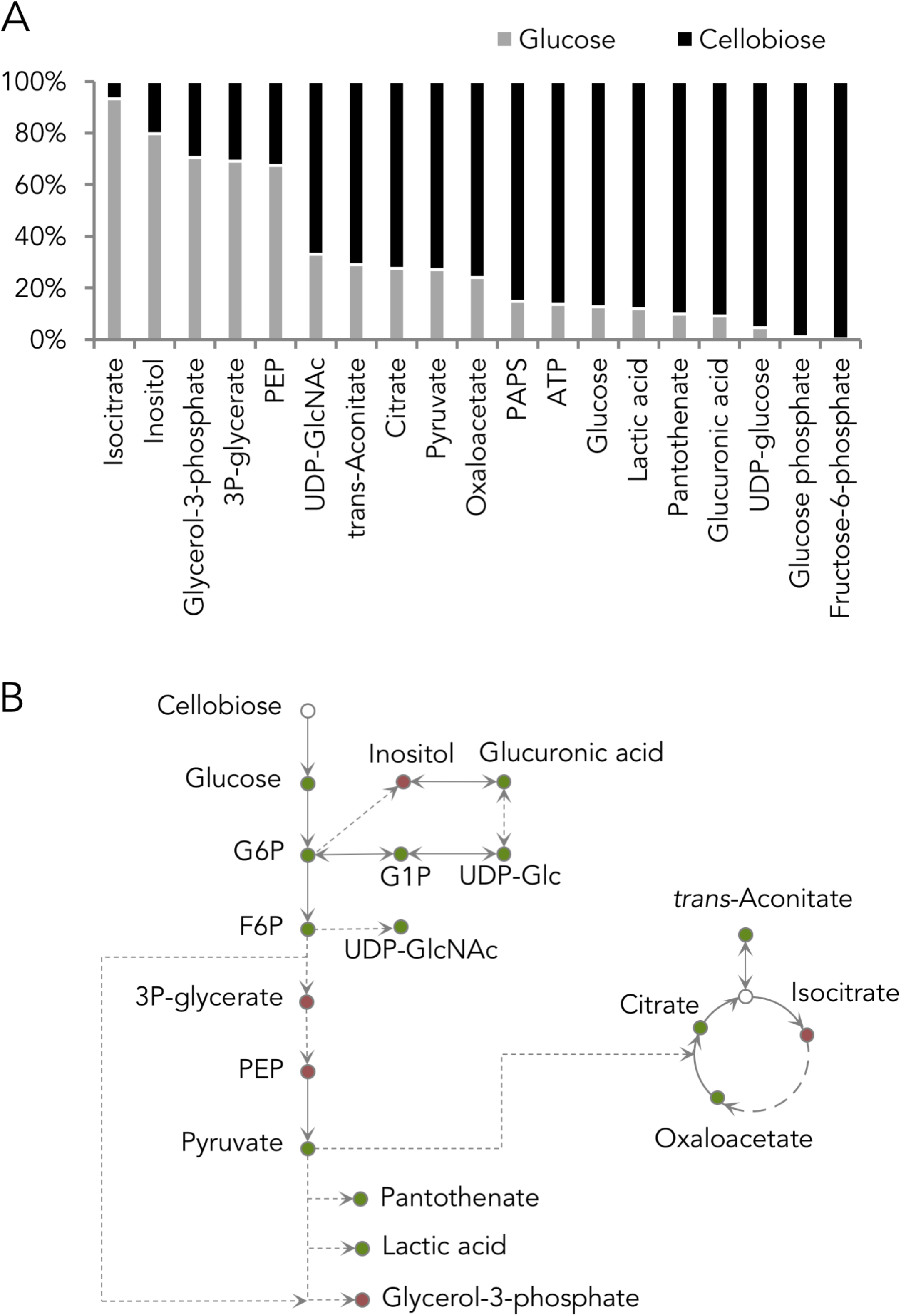
Metabolite profile of cells provided with glucose or cellobiose. (A) Significant changes of intracellular metabolite levels in cells provided with cellobiose compared to cells provided glucose as a sole carbon source. (B) Schematic representation of metabolite changes. The green and red dots represented higher and lower relative metabolite levels in cells provided with cellobiose compared to cells provided with glucose. Statistical analyses are described in the Materials and Methods.

### Phosphofructokinase-1 inhibition by excess ATP

Given the dramatic buildup of glucose, G6P and F6P intermediates prior to the phosphofructokinase (*PFK1*) reaction in glycolysis (Figure 1B, 2A), we reasoned that Pfk1 might be a major bottleneck in cellobiose consumption. Pfk1 catalyzes the phosphorylation of F6P, using one ATP and yielding fructose 1,6-bisphosphate (F1,6BP) as a product. As the second committed step in glycolysis, Pfk1 is heavily regulated—with ATP acting as an allosteric inhibitor and AMP and fructose 2,6-bisphosphate serving as activators (8–10).

**Figure 2.**
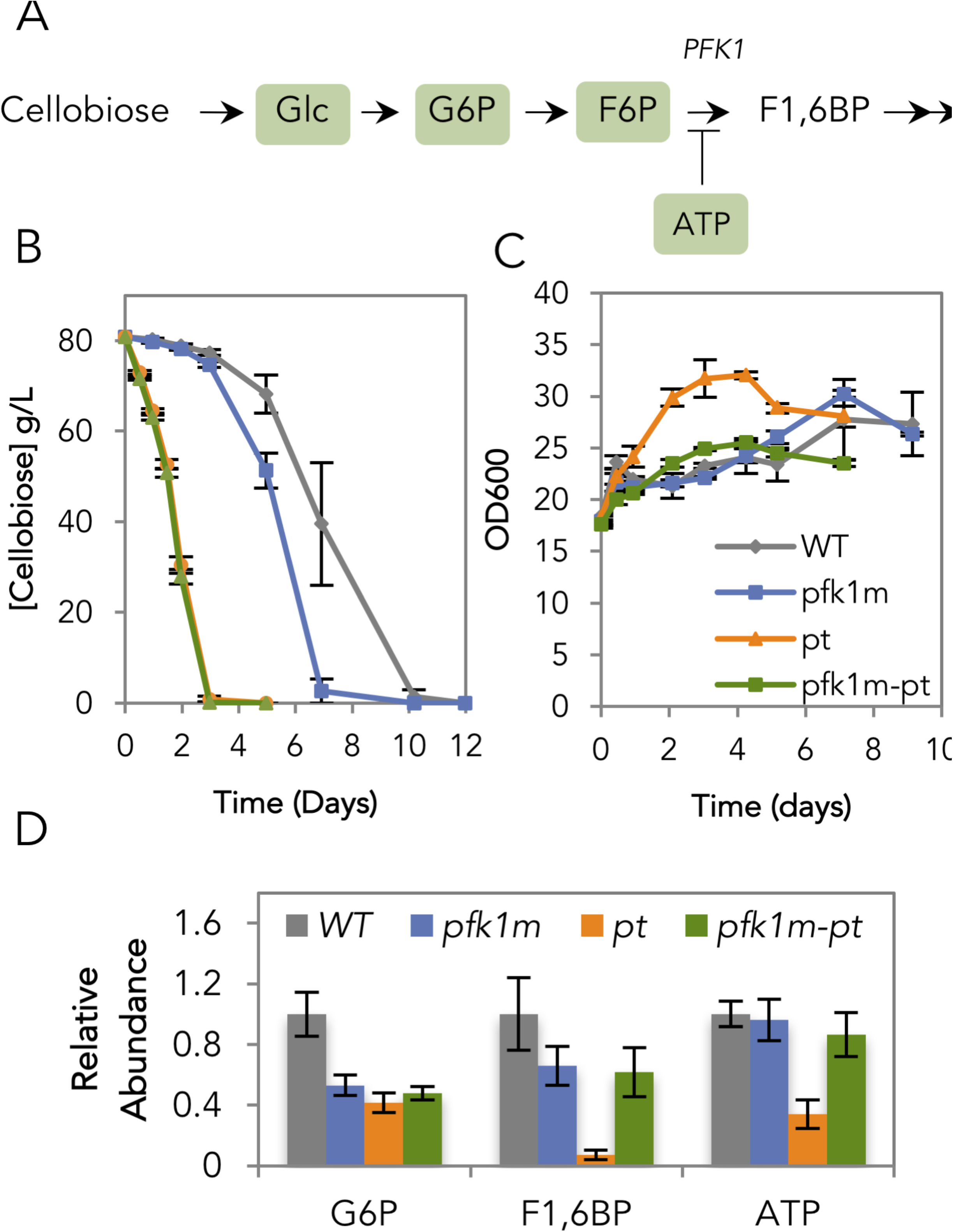
Manipulation of phosphofructokinase (*PFK1*) and plasma membrane ATPase *(PMA1*). (A) Schematic representation of cellobiose consumption route in the upper glycolytic pathway. Glc, G6P, F6P and ATP are highlighted, and were found in higher abundance when cellobiose was provided in comparison to glucose. (B) Cellobiose consumption profile and (C) cell density profile of the strains with ATP-insensitive Pfk1 (*pfk1m*), constitutively active Pma1 (*pt*) and the combination of both mutations (*pfk1m-pt*) in comparison to the cellobiose pathway-only strain, here used as wild-type (WT). (D) Relative abundance of G6P, F1,6BP and ATP levels of the WT, *pfk1m, pt* and *pfk1m-pt* strains, relative to the WT strain fermenting cellobiose. The experiments were carried out in 5 biological replicates, with standard errors from the mean shown.

To test whether allosteric inhibition of Pfk1 by ATP limited cellobiose fermentation, a mutation in Pfk1 that makes the enzyme ATP-insensitive (P722L in the Pfk1 beta subunit, note the original publication referred to this mutation as P728L (11)) was introduced into the chromosomally encoded *PFK1* gene (mutation here termed *pfk1m*) in the cellobiose utilizing strain. This mutation was previously shown to reduce ATP inhibition but also AMP and fructose 2,6-bisphosphate activation of Pfk1 in *S. cerevisiae* (11). We chose this mutation over an ATP-insensitive, AMP/F2,6BP-sensitive mutant phosphofructokinase (10) because the latter's phenotype has not been evaluated in *S. cerevisiae*. High initial cell densities were used hereafter, as the focus of this study is sugar consumption rather than growth.

Consistent with allosteric inhibition of Pfk1 by ATP, the cellobiose consumption efficiency (E_c_) of the *pfk1m* strain increased by 33% in comparison to the strain with wild-type Pfk1 (Figure 2B). In these high cell densities, negligible changes in growth rate were observed (Figure 2C). The relative abundance of G6P and F1,6BP decreased by 47% and 34%, respectively, while that of ATP remained relatively unchanged (Figure 2D). The unchanged ATP level was expected as the ATP requirement for the cellobiose consumption pathway was likely offset by the ATP generated as part of carbon metabolism. These results indicate that the 6-fold increase in cellular ATP concentrations allosterically inhibited Pfk1, resulting in accumulation of glucose, G6P and F6P, which eventually slowed down cellobiose consumption.

### Limited activity of plasma membrane ATPase

Although the *pfk1m* strain partially increased the rate of cellobiose fermentation, cellular ATP remained elevated relative to glucose fermentation. It is unlikely that ATP production was the cause of the difference, as fermentation is limited to substrate-level phosphorylation under anaerobic conditions regardless of carbon source. We therefore tested whether the activity of one of the major ATP sinks in yeast, the plasma membrane ATPase (Pma1) was responsible for the ATP buildup. Pma1 hydrolyzes 25-40% of cellular ATP in yeast (12) and is heavily regulated by glucose (13).

A constitutively active mutant form of *PMA1* (*pma1*-Δ*916*, here abbreviated *pt*) (14) was introduced into the endogenous *PMA1* locus in the cellobiose utilizing strain. This mutation results in high Pma1 ATPase activity even in carbon starvation conditions (14). The E_c_ of the *pt* strain was 4 times that of the control (Figure 2B), whereas the growth rate of *pt* strain was only slightly faster than the Pma1 WT strain (Figure 2C). As expected, we observed a 66% decrease in cellular ATP levels in the *pt* strain in comparison to the wild-type control (WT, i.e. cellobiose pathway only) (Figure 2D). In addition, the concentrations of G6P and F1,6BP decreased by 58% and 93%, respectively, relative to strains expressing wild-type *PMA1*. Notably, these concentrations dropped more than when the ATP-insensitive *PFK1* mutant was introduced (Figure 2D). These results suggest that increased Pma1 ATPase activity improved cellobiose fermentation. We hypothesize that the drastic decrease in F1,6BP level and the fast growth rate were the result of rapid glycolytic flux, as the cells experience low cellular ATP levels in the *pt* strain.

Next, we observed the phenotypes of *pfk1m-pt* double mutant strain. The cellobiose consumption profile of a *pfk1m-pt* double mutant was identical to that of the *pt* strain (Figure 2A). However, the growth rate and relative abundance of G6P, F1,6BP and ATP of the *pfk1m-pt* differed from the *pt* strain (Figure 2C,D). In fact, their levels were similar to those in the *pfk1m* strain. These results imply that while the ATP might be hydrolyzed rapidly due to the *pt* effect, the removal of ATP inhibition on *pfk1* allowed enough ATP to be regenerated downstream that no growth burst was observed. The underlying explanation of the mixed phenotypes will require future experiments to dissect how Pfk1 exerts allosteric control on glycolysis and ATP levels.

### Carbon starvation-like state of the plasma membrane ATPase

Although cellobiose theoretically provides the same energy and carbon availability to cells as glucose, its releases glucose only after intracellular hydrolysis by β-glucosidase. Thus the cellobiose consumption system used here does not generate extracellular glucose, which acts as a crucial signaling molecule for yeast carbon metabolism. Taken together with the observation that increased ATPase activity in the *pt* strain increased cellobiose consumption efficiency, we wondered whether the limited Pma1 activity in cellobiose-fed cells is due to the absence of extracellular glucose in the media. Transcriptionally, the presence of glucose increases *PMA1* mRNA levels by 2-4 times via the regulation of Rap1, Gcr1 and Sir2 (15–17). Consistent with the requirement for extracellular glucose sensing, previous RNA sequencing experiments revealed a 40% decrease in *PMA1* transcript levels when cellobiose was provided in place of glucose (5). However, although transcriptional regulation of *PMA1* is important, it is slower than post-transcriptional regulation and results in smaller changes (13, 18).

In the presence of glucose, phosphorylation of Ser-899 decreases Pma1's K_m_ and Ser-911/Thr-912 increases Pma1's V_max_ for ATP, respectively (13, 19, 20). Given the 6-fold excess amount of ATP observed in cellobiose utilizing conditions (Figure 1A), the effective velocity of the Pma1 reaction is likely approaching V_max_ regardless of the phosphorylation status at Ser-899 (21). The K_m_ of ATP hydrolysis by Pma1 has been reported to increase approximately 3-fold from 1.2 mM in glucose-fermenting cells to 4.0 mM in glucose-starved cells, while a 10-fold decrease in V_max_ was reported in the same study (22). We reasoned that the 6-fold increase of ATP in cellobiose-fed cells should result in ATP concentrations in excess of the K_m_ for Pma1, resulting in Pma1 activity being limited by its V_max_. Thus, in this study, we did not investigate the phosphorylation of Ser-899 and chose to investigate whether V_max_-determining phosphorylation states of Ser-911 and Thr-912 might play a major role in establishing the efficiency of cellobiose fermentation (Figure 3A).

**Figure 3.**
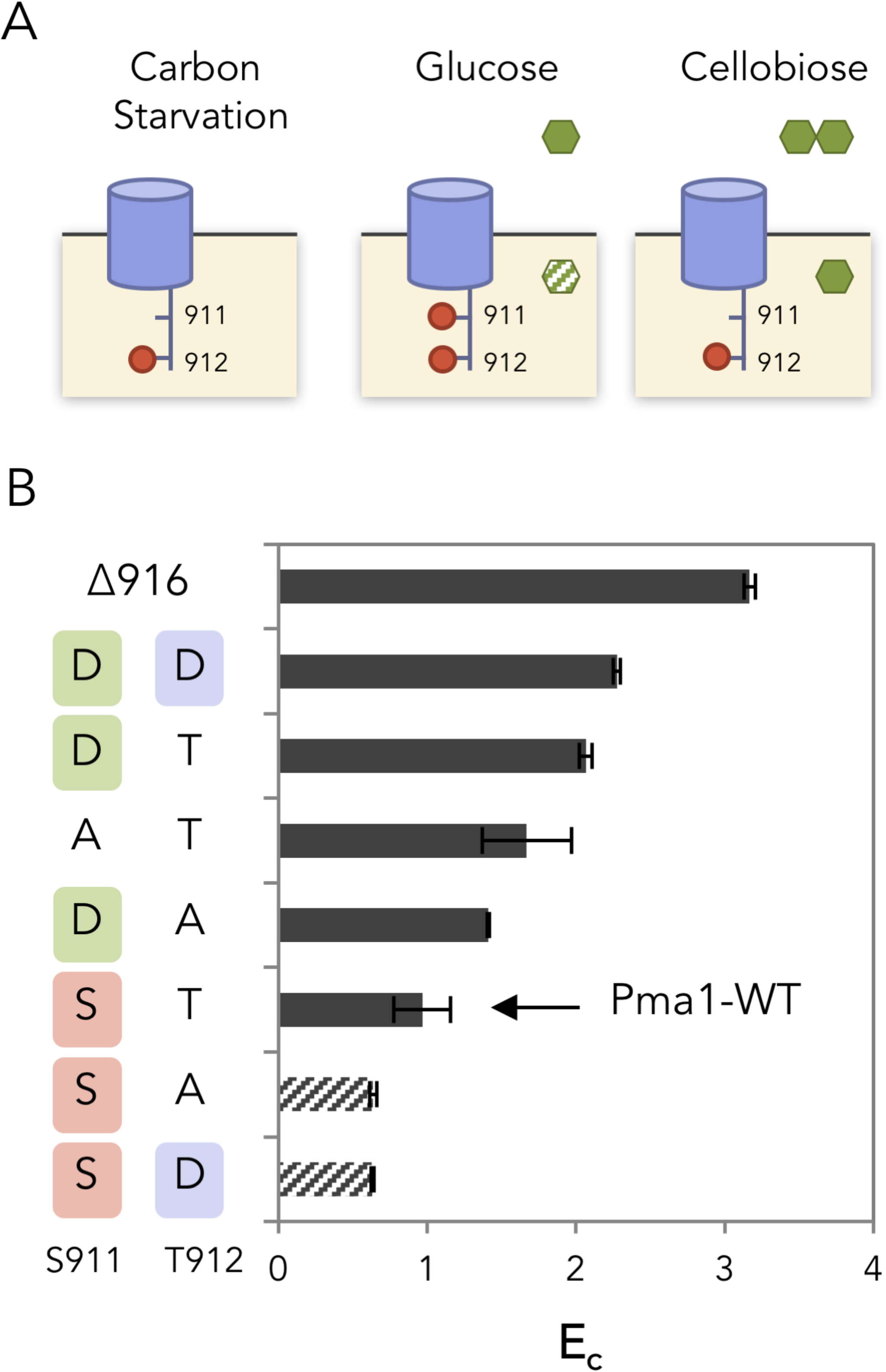
Carbon starvation-like state of the plasma membrane ATPase (*PMA1*) in cellobiose-fermenting cells. (A) Phosphorylation states of Pma1 residues S911 and T912 under carbon starvation, glucose metabolizing and cellobiose metabolizing conditions. (B) Cellobiose consumption efficiency (E_c_) of cells expressing Pma1 with phosho-mimic or phosphorylation-preventing mutations at positions serine 911 (S911) and threonine 912 (T912). Shown are the mean and standard deviation for 3 biological replicates. The experiments were carried out in 2 biological replicates, with standard errors from the mean shown.

Combinatorial mutations of Ser-911/Thr-912 to alanine and aspartic acid were introduced into the endogenous *PMA1* gene to prevent or mimic phosphorylation, respectively. We were unable to obtain strains with *pma1-*S911A/T912A and *pma1-* S911A/T912D, potentially because the combinations were lethal. All mutant strains whose Pma1 S911 position was mutated to aspartic acid consumed cellobiose more efficiently in comparison to when the S911 position remained unchanged (Figure 3B). By contrast, mutating Pma1 T912 to aspartic acid did not show a correlation with the cellobiose consumption phenotype. These results suggest that phosphorylation of Pma1 at S911 was lacking when cellobiose was provided as a sole carbon source.

### Positive effects of extracellular glucose sensor deletions

According to the above mutational analysis, the Pma1 phosphorylation state of cellobiose-fed cells was similar to that in carbon starvation conditions (Figure 3A) (20). In the previously published experiments (20), carbon-starved cells were prepared by incubating mid-exponential phase cells in media without glucose. In such conditions, neither extracellular nor intracellular glucose is present. For the cellobiose-fed cells, based on the relatively high level of intracellular glucose we detected (Figure 1A), it is unlikely that the intracellular glucose induced Pma1 carbon starvation. Additionally, since intracellular glucose metabolism is expected in cellobiose-fed cells, its effect on Pma1 carbon starvation was also ruled out (23). We therefore tested the role of extracellular glucose in regulating Pma1 activity. In cellobiose-fed cells, glucose is not provided as part of the media, and thus the extracellular glucose is absent. We hypothesize that the extracellular glucose is likely essential for full activation of Pma1 through S911 phosphorylation.

Snf3, Rgt2 and Gpr1 are the three known sensors of extracellular glucose in *S. cerevisiae*. Snf3 and Rgt2 mainly regulate glucose transport while Gpr1 controls cell physiology via an interaction with Gpa2 to activate protein kinase A and cAMP synthesis (23). To mimic the presence of extracellular glucose, constitutively active mutations (*snf3* R229K*, rgt2* R231K and *gpa2* R273A) were introduced into the endogenous loci to probe the role of each glucose-sensing pathway (24, 25). Surprisingly, the cellobiose consumption efficiency of all the three mutant strains decreased by ∼25% (Figure S2A). We then inverted the genetic modifications by deleting *SNF3, RGT2* and/or *GPA2*. Notably, the triple glucose-sensing deletion strain (*snf3*Δ*rgt2*Δ*gpa2*Δ*, or srg*Δ) showed a 275% increase in E_c_ (Figure 4A).

**Figure 4.**
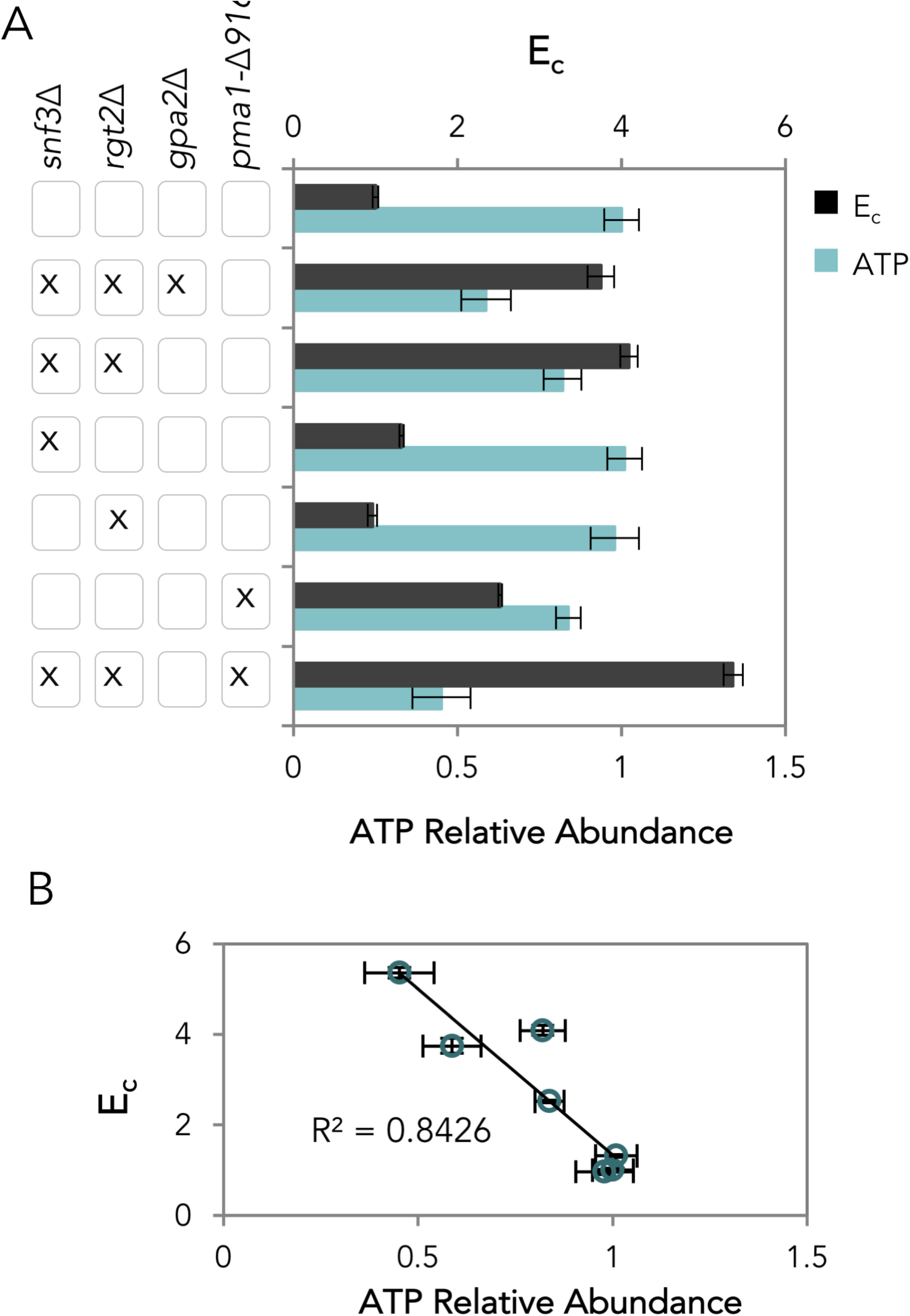
Effect of glucose sensor deletions. (A) Cellobiose consumption efficiency (E_c_) and cellular ATP levels of strains with different glucose sensor deletions and different combinations of constitutively-active Pma1 mutations. Shown are the averages and standard deviations of three biological replicates. (B) Correlation of E_c_ and cellular ATP. Standard deviations for three biological replicates are shown for each point.

Combinatorial deletions revealed that the Gpr1 pathway did not contribute to improved cellobiose fermentation, but combining the *SNF3* and *RGT2* deletions (*sr*Δ) was necessary and sufficient to replicate the E_c_ of the triple deletion strain (Figure 4A, Figure S2B, C). Consistent with the observed E_c_ values, the intracellular ATP levels of *srg*Δ and *sr*Δ decreased by 41% and 18% respectively, while those in the individual-deletion strains *snf3*Δ and *rgt2*Δ remained unchanged (Figure 4A). These results reveal a negative correlation between E_c_ and cellular ATP levels (Figure 4B) and that Snf3 and Rgt2 acted together to regulate cellular ATP levels, in addition to regulating glucose transport.

Although the additional deletion of *GPA2* (*gpa2*Δ) in the *sr*Δ strain did not further improve E_c_ (Fig 4A), it reduced the relative abundance of ATP by 28%, implying that the Gpr1 pathway has a separate mechanism to control cellular ATP levels that does not directly affect carbon metabolism. Consistently, *gpa2*Δ had a negative or neutral impact on cellobiose consumption (Figure S2). The decrease in ATP level may be a result of altered cellular activities, controlled by Gpr1 via the Tor and cAMP-PKA-Ras pathways (23). The relationship between Gpr1-regulated ATP levels and carbon metabolism remains to be discovered. Since *gpa2*Δ did not have a direct effect on E_c_, it was not investigated further in this study.

### Snf3/Rgt2 regulation of cellular ATP levels

To examine whether Snf3/Rgt2 regulated the cellular ATP level in cellobiose fermentations via Pma1, an *snf3*Δ *rgt2*Δ *pma1-*Δ*916* strain (*sr*Δ*-pt*) was constructed. Notably, the E_c_ of the *sr*Δ*-pt* strain increased more than 4 times in comparison to the wild-type control (Figure 4A, Figure 5A). The improvement was additive, within the range of the ΔEc summation of *sr*Δ and *pma1-*Δ*916* strains relative to wild-type (Figure 5A). Although ATP levels decreased in a nearly linear fashion as a function of E_c_ (Figure 4B), it is not presently possible to ascertain whether the *sr*Δ and *pma1-*Δ*916* mutations act entirely independently due to limitations in measurement accuracy (Figure 5A).

**Figure 5.**
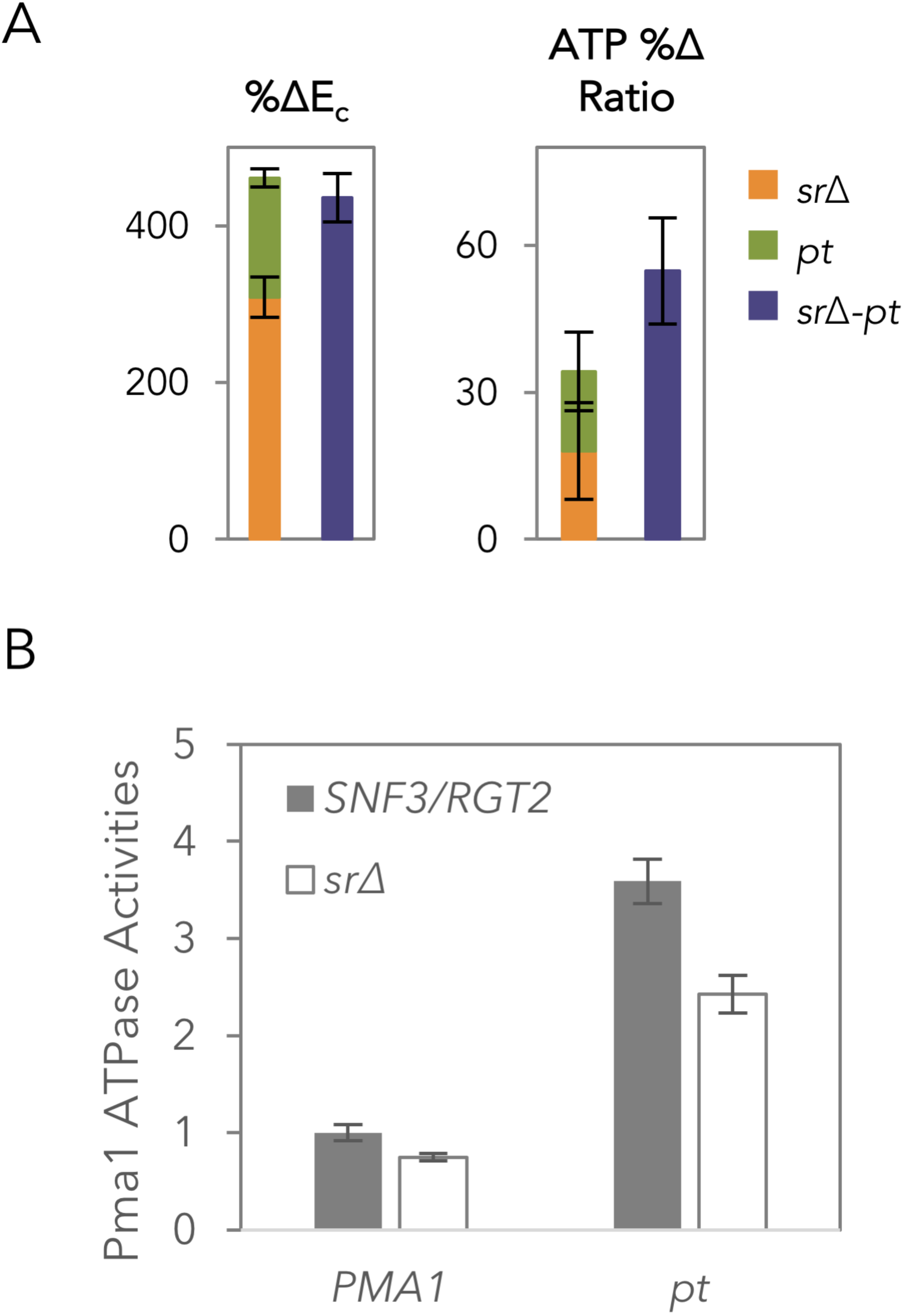
Glucose sensor deletions and cellular ATP levels. (A) Additive effect of the increase in E_c_ and the decrease in cellular ATP of *sr*Δ and *pt* in *sr*Δ*-pt* strain. The experiments were carried out in 5 biological replicates, with standard errors form the mean shown. (B) Specific Pma1 ATPase of WT*, pt, sr*Δ and *sr*Δ*-pt* strains measured from normalized membrane fractions of cells harvested at mid-log phase. The experiments were carried out in 3 biological replicates, with standard errors from the mean shown.

To further determine the relationship between Snf3/Rgt2 and Pma1, the vanadate-specific ATPase activity of Pma1 (26) from different strains consuming cellobiose was analyzed (Figure 5B). Consistent with the constitutively active nature of the *pt* mutation, the activities of Pma1 in the *pt* and *sr*Δ*-pt* strains were higher than those observed in the WT or *sr*Δ strains, respectively. Addition of *sr*Δ decreased the Pma1 ATPase activities by 25% and 32% in WT and *pt* strains, respectively. In other words, the absence of Snf3 and Rgt2 led to a partial decrease in Pma1 ATPase activity, which implied that Snf3/Rgt2 partially activated Pma1 ATPase activity in the absence of glucose.

## Discussion

To identify the effects of a minimal alteration to carbon metabolism in yeast, we chose a cellobiose-consumption pathway composed of two genes and analyzed its cellular metabolite profiles in comparison to cells provided with glucose, yeast's preferred carbon source (Figure 6). Here we focus on the cellobiose consumption efficiency (E_c_), as E_c_ linearly correlated with ethanol production rate, while ethanol yield remained mostly unchanged (Figure S3).

**Figure 6.**
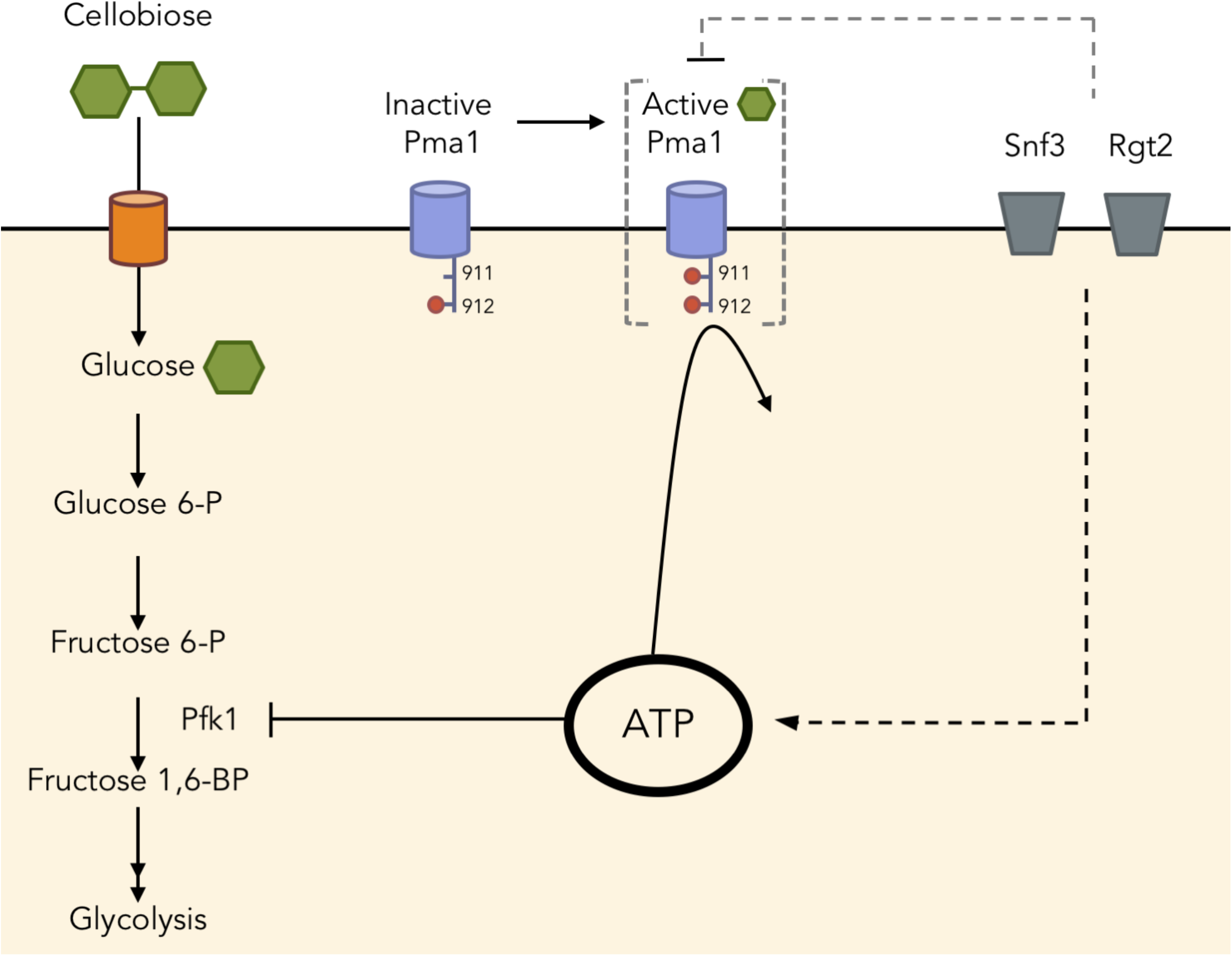
Schematic representation of ATP homeostasis and cellular regulation of cellobiose fermentation. Excess ATP inhibited phosphofructokinase (Pfk1), resulting in an upper glycolytic metabolite buildup and slow cellobiose consumption. The buildup of ATP may be caused by low activity of Pma1, with no phosphorylation at position 911. Additionally, glucose sensors Snf3 and Rgt2 together influenced cellular ATP levels via Pma1 and other mechanisms to be identified.

More than half of the metabolites significantly changed in abundance when cellobiose was provided in place of glucose. The buildup of G6P, F6P and ATP in *S. cerevisiae* fermenting cellobiose suggested that Pfk1 was one of the bottlenecks in the process. Pfk1 is subjected to complex allosteric regulation, including inhibition by ATP and activation by AMP and fructose 2,6-bisphosphate (F2,6BP) (8–10). The Pfk1 bottleneck was partially relieved in cells expressing an ATP/AMP/F2,6BP-insensitive *PFK1* allele, while the ATP level remained elevated. These results contrast with previous studies that identified ATP depletion and the buildup of fructose-1,6-bisphosphate–the metabolite immediately downstream of Pfk1–as a weak link in glycolysis (1). Although we did not investigate the effect of AMP and F2,6BP activation since their changes between glucose and cellobiose conditions were less than 2 fold and they did not meet the significance threshold of a p-value less than 0.01, it is possible that they could influence cellobiose consumption efficiency.

ATP is central to a cell's energy currency, but too much ATP is not necessary beneficial (27, 28). In fact, we observed a negative correlation between cellular ATP levels and cellobiose consumption efficiency (Figure 4B). A similar correlation has been reported for glucose as a carbon source, suggesting metabolic uncoupling of energy homeostasis in yeast cells (29). We propose that intracellular glucose concentrations– generated by cellobiose hydrolysis in our experiments–and glucose metabolism (23) are insufficient to trigger glucose activation of key metabolic pathways and enzyme activity. For example, we found that the ATP-dependent proton pump Pma1 existed in a carbon-starvation like state during cellobiose fermentation, and was partially responsible for the aberrant accumulation of ATP. These results suggest that neither intracellular glucose nor glucose metabolism are sufficient to fully activate Pma1. A previous study showed the lack of phosphorylation of S899 and S911/T912 in Pma1, in a hexokinase/glucokinase deletion strain (*hxk1*Δ *hxk2*Δ *glk1*Δ) provided with glucose, suggesting that phosphorylation of these residues requires glucose metabolism (30). Together with our results, we propose that the activation of Pma1 through S911 phosphorylation requires both extracellular glucose and glucose metabolism. Our results reveal that the cellobiose utilization system allows uncoupling of glucose metabolism and intracellular glucose from extracellular glucose signaling. Future experiments will be required to reveal why ATP was not consumed by other cellular processes triggered under starvation (31).

Cytosolic pH is also a key regulator of carbon utilization (32), and is likely to be impacted by the use of the proton symporter CDT-1 for cellobiose import and the resulting low activity of Pma1. High cytosolic pH is necessary and sufficient to activate Tor-Ras-PKA activities, which are downstream of Gpr1 glucose sensing pathway (32). By contrast, the proton symporter CDT-1 and low activity of Pma1 may result in a low cytosolic pH. However, cytosolic pH alone is unlikely to determine the cellobiose consumption efficiency (E_c_) as the strain with an ATP/AMP/F2,6BP-insensitive *PFK1* allele showed improved Ec but unaltered cellular ATP levels. Furthermore, the inactivation of the Gpr1 pathway resulted in decreased cellular ATP but unaltered E_c_. Glucose storage, i.e. in the form of trehalose, may be interconnected through the Gpr1 pathway because Ras-cAMP activates trehalase required to break down trehalose—a phenomenon observed when gluconeogenesis is switched to glycolysis (33). Trehalose cycling has been shown to lead to an imbalanced state of glycolysis (1). The relationship between carbon storage and E_c_ will require future studies to examine this relationship.

We also found that the well-studied extracellular glucose sensors Snf3/Rgt2 exhibited a novel role in cellular ATP homeostasis partially through the major plasma membrane ATPase Pma1. Deletion of extracellular glucose sensors (Snf3/Rgt2) increased cellobiose consumption efficiency and partially restored ATP levels. Interestingly, the absence of the Snf3/Rgt2 decreased Pma1 ATPase activities, an effect that should have led to an increase in ATP level. The restored low ATP level observed in the *sr*Δ strain implied that Snf3/Rgt2 regulated cellular ATP level with additional mechanism(s) other than through Pma1. It is known that deletion of *SNF3* and *RGT2* slows down glucose consumption (34), due to the inability of these strains to degrade Mth1/Std1, which block the promoter regions of hexose transporters required for optimal glucose import (35–38). Unlike glucose, cellobiose does not signal Mth1 degradation even with intact Snf3/Rgt2 (38). Thus, genes downstream of Mth1 regulation, including hexose transporters, are not expected to be responsible for the improved E_c_ and decreased cellular ATP levels observed in *sr*Δ. Consistent with this model, no growth defect is observed in a *mth1*Δ strain growing on glucose, suggesting that Snf3/Rgt2 has additional regulatory nodes other than Mth1 (34). Future transcriptional profiling and ribosome profiling experiments will be required to reveal the additional Snf3/Rgt2 roles in cellular ATP homeostasis.

The present systems-level study of a minimal synthetic biology pathway for cellobiose consumption revealed the dramatic impact of decoupling extracellular and intracellular glucose sensing, resulting in an overabundance of ATP in cells. The inability of *S. cerevisiae* to catabolize ATP for cellular processes in the presence of intracellular glucose and glucose metabolism but in the absence of extracellular glucose resulted in slow fermentation. Thus, ATP levels must be kept in a relatively narrow range for optimal fermentation and to allow robust startup of glycolysis, yet yeast seems to lack a direct mechanism to monitor ATP concentrations. For example, a dynamic model showed that a small concentration difference of inorganic phosphate, a product of ATP hydrolysis, could alter cell fate from a stable glycolytic steady state to an imbalanced dead-end state (1). Here, we found that the extracellular glucose sensing by Snf3/Rgt2 required for optimal glucose fermentation (2) can be uncoupled from the role of these receptors in regulating ATP homeostasis under carbon starvation conditions. It will be important in the future to map the fully regulatory pathways of ATP homeostasis leading from Snf3/Rgt2 and, independently, terminating in Pma1.

## Materials & Methods

### Yeast strains, media and anaerobic fermentation

The *S. cerevisiae* background strain used in this study was S288C *ura3::P_PGK1_-cdt-1* N209S/F262Y*-T_ADH1_lyp1::P_TDH3_-gh1-1 (codon optimized)-T_CYC1_* derived by chromosomal DNA library selection (39). The strain was subjected to further modifications using the CRISPRm system (39). The list of strains constructed, CRISPRm guides and primers used are included in Table S1.

Seed cultures for cellobiose fermentation experiments were grown in 20 g/L glucose modified optimal minimal media (oMM) (5) and harvested at mid-log phase. All cellobiose fermentation experiments were conducted under strict anaerobic conditions, in 80 g/L cellobiose oMM media at an initial OD_600_ of 20, using 10 mL serum flasks containing 5 mL fermentation in 2-5 biological replicates. The flasks were incubated at 30°C, 220 rpm. The cellobiose consumption efficiency (E_c_) was defined as the inverse of the area under the curve of extracellular cellobiose concentration over time.

### Analytical analysis of yeast metabolites

Extracellular cellobiose concentrations were determined by high performance liquid chromatography on a Prominence HPLC (Shimadzu, Kyoto, Japan) equipped with Rezex RFQ-FastAcid H 10 × 7.8 mm column. The column was eluted with 0.01 N of H_2_SO_4_ at a flow rate of 1 mL/min, 55°C.

For the metabolite profiling comparison between glucose and cellobiose (modified from (40)), equal amounts of yeast cells at mid-exponential phase of anaerobic sugar consumption (10 g/L cellobiose or glucose) were harvested (final pellet OD_600_ equivalent to 5). The samples were quenched in 180 µL of 40:40:20 acetonitrile:methanol:water. Following the addition of 10 nmols of d3 serine (as an internal standard), the mixtures were vortexed and centrifuged at 13,000 rpm for 10 minutes. The supernatants were injected onto an Agilent 6460 QQQ LC-MS/MS and the chromatography was achieved by normal phase separation with a Luna NH_2_ column (Phenomenex) starting with 100% acetonitrile with a gradient to 100% 95:5 water acetonitrile. 0.1% formic acid or 0.2% ammonium hydroxide with 50 mM ammonium acetate was added to assist with ionization in positive and negative ionization mode, respectively. Five biological replicates were used for each sample analyzed.

For targeted intracellular metabolite comparisons, yeast cells equivalent to 20 OD_600_ units were harvested and filtered through a 0.8 µm nylon membrane, prewashed with 3mL water, followed by another 3 mL water wash after cell filtration. The membranes were placed in 1.5mL extraction solution (0.1 M formic acid, 15.3 M acetonitrile) flash-frozen in liquid nitrogen, and stored at −80°C. Before analysis, the extracts were vortexed for 15 minutes and centrifuged to collect the supernatants at 4°C. Glucose 6-phosphate and fructose 1,6-bisphosphate were separated and identified using a 1200 Series liquid chromatography instrument (Agilent Technologies, Santa Clara, CA). 1 µL of each sample was injected onto an Agilent Eclipse XDB-C18 (2.1 mm i.d., 150 mm length, 3.5 µm particle size) column with a Zorbax SB-C8 (2.1 mm i.d., 12.5 mm length, 5 µm particle size) guard column and eluted at 25 °C and a flow rate of 0.2 mL/min with the following gradient (modification from (41)): 15 min isocratic 100% buffer A (10 mM tributylamine/15 mM acetic acid), then in 15 min with a linear gradient to 60% buffer B (methanol), 2 min isocratic 60% B, then 10 min equilibration with 100% buffer A. The eluent from the column was introduced into a mass spectrometer for 25 minutes after the first 10 minutes. Mass spectrometry (MS) was performed on an LTQ XL ion trap instrument (Thermo Fisher Scientific, San Jose, CA) with an ESI source operated in negative ion mode. The MS settings were capillary temperature 350 °C, ion spray voltage 4.5 kV, sheath gas flow: 60 (arbitrary units), auxiliary gas flow 10 (arbitrary units), sweep gas flow 5 (arbitrary units). For the MS/MS product ion scan, the scan range was m/z 80 to m/z 300. The compounds G6P at m/z 259.1 and F16BP at m/z 339.1 were isolated with an m/z 2 isolation width and fragmented with a normalized collision-induced dissociation energy setting of 35% and with an activation time of 30ms and an activation Q of 0.250.

The significance threshold between cells provided with cellobiose and cells provided with glucose was set at a p-value of 0.01. Five biological replicates were used in each sample group. Only the metabolites with higher than a 2 fold-change between each sample group were included in the analysis.

### Plasma membrane isolation

Strains were subjected to cellobiose fermentation under anaerobic conditions. Yeast cells with an OD_600_ equivalent to 40 were harvested at mid-log phase and flash frozen in liquid nitrogen. Membrane fractions were extracted based on the protocol published in (42).

### Pma1 ATPase activity assay

The ATPase assay described in (26) was modified as follows. 30 µg of the isolated membrane fraction was incubated in assay buffer (50 mM MES pH 5.7, 10 mM MgSO4, 50 mM KCl, 5 mM NaN_3_, 50 mM KNO_3_) with and without 3 mM orthovanadate for 25 minutes at 30 °C. 1.8 mM ATP was added to start the 100 uL reactions. The reactions were incubated at 30 °C for 15 minutes, then the membranes were isolated from the reactions by centrifugation at 13,000x g for 10 minutes at 4°C. The released inorganic phosphate was measured in the supernatant using the ATPase/GTPase Activity Assay Kit (Sigma-Aldrich) based on the manufacturer's protocol. The specific Pma1 ATPase activities were calculated by subtracting the concentration of released inorganic phosphate in reactions provided with orthovanadate from those without.

### Yeast cell-based cellobiose uptake assay

The cell-based cellobiose uptake assay was modified from (43). Yeast strains were grown to mid-exponential phase in 2% oMM glucose, washed with assay buffer (5 mM MES, 100 mM NaCl, pH 6.0) three times and resuspended to a final OD_600_ of 10. Equal volumes of the cell suspension and 200 µM cellobiose were mixed to start the reactions, which were incubated at 30°C with continuous shaking for 15 minutes. The reactions were stopped by adding 150 µL of supernatants to 150 µL 0.1 M NaOH. The concentrations of the remaining cellobiose were measured using an ICS-3000 Ion Chromatography System (Dionex, Sunnyvale, CA, USA) equipped with a CarboPac^®^ PA200 carbohydrate column. The column was eluted with a NaOAc gradient in 100 mM NaOH at a flow rate of 0.4 mL/min, 30°C.

## Acknowledgements

The authors thank Raissa Estrela, Dr. Xin Li, Dr. Ligia Acosta-Sampson, Dr. Yuping Lin and Dr. Matt Shurtleff for helpful discussions. This work was supported by funding from Energy Biosciences Institute to JHDC and from National Institutes of Health (R01CA172667) to DKN.

## Conflict of Interest

The authors declare that they have no conflict of interest.

## Author contributions

All authors contributed to the design of the experiments. KC and DIB carried out the experiments. DKN and JHDC aided in interpretation of the systems-level experiments. KC wrote the manuscript, with editing from DIB, DKN, and JDHC.

## Supplementary Figure legends

**Figure S1. Metabolite profile of cells provided with glucose or cellobiose.** (A) cell density and sugar consumption profiles of the strains used for metabolite profiling experiments. (B) Heatmap representation of steady state intracellular metabolite abundance of cells provided with glucose or cellobiose under anaerobic conditions. Shown are the results of 5 biological replicates. Asterisks (*) mark identified compounds that were significantly different between glucose and cellobiose conditions at a significance threshold of a p-value < 0.01.

**Figure S2. Cellobiose consumption profiles and efficiency of Snf3, Rgt2 and Gpa2 mutants.** (A) Cellobiose consumption profiles of strains with consitutively active Snf3 (R229K), Rgt2 (R231K) and Gpa2 (R273A). (B) Cellobiose consumption profiles of *SNF3*, *RGT2*, and *GPA2* double deletion combinations. (C) Cellobiose consumption profiles of *SNF3* and *RGT2* single deletion in comparison to the *sr*Δ*, srg*Δ *and sr*Δ*-pt* strains. (D) Cellobiose consumption profiles of *GPA2* deletion (*gΔ)* in comparison to the WT strain. In panels A, B, C, and D, the results of 3 biological replicates are shown.

**Figure S3. Relationship of ethanol productivity parameters and cellobiose consumption efficiency (E_c_).** Ethanol production rate and the final concentration of ethanol are plotted against E_c_.

**Table S1: List of strains constructed, CRISPRm guides and primers.**

## References

1. van Heerden JH, Wortel MT, Bruggeman FJ, Heijnen JJ, Bollen YJM, Planque R, Hulshof J, O'Toole TG, Wahl SA, Teusink B. 2014. Lost in Transition: Start-Up of Glycolysis Yields Subpopulations of Nongrowing Cells. Science 343:1245114–1245114.

2. Youk H, van Oudenaarden A. 2009. Growth landscape formed by perception and import of glucose in yeast. Nature 462:875–879.

3. Ha S-J, Galazka JM, Kim SR, Choi J-H, Yang X, Seo J-H, Glass NL, Cate JHD, Jin Y-S. 2011. Engineered Saccharomyces cerevisiae capable of simultaneous cellobiose and xylose fermentation. Proc Natl Acad Sci USA 108:504–509.

4. Galazka JM, Tian C, Beeson WT, Martinez B, Glass NL, Cate JHD. 2010. Cellodextrin transport in yeast for improved biofuel production. Science 330:84– 86.

5. Lin Y, Chomvong K, Acosta-Sampson L, Estrela R, Galazka JM, Kim SR, Jin Y-S, Cate JH. 2014. Leveraging transcription factors to speed cellobiose fermentation by Saccharomyces cerevisiae. Biotechnology for Biofuels 7:126.

6. Daran-Lapujade P, Rossell S, van Gulik WM, Luttik MAH, de Groot MJL, Slijper M, Heck AJR, Daran J-M, de Winde JH, Westerhoff HV, Pronk JT, Bakker BM. 2007. The fluxes through glycolytic enzymes in Saccharomyces cerevisiae are predominantly regulated at posttranscriptional levels. Proceedings of the National Academy of Sciences 104:15753–15758.

7. Kim H, Lee W-H, Galazka JM, Cate JHD, Jin Y-S. 2014. Analysis of cellodextrin transporters from Neurospora crassa in Saccharomyces cerevisiae for cellobiose fermentation. Appl Microbiol Biotechnol 98:1087–1094.

8. Bañuelos M, Gancedo C, Gancedo JM. 1977. Activation by phosphate of yeast phosphofructokinase. J Biol Chem 252:6394–6398.

9. Avigad G. 1981. Stimulation of yeast phosphofructokinase activity by fructose 2,6-bisphosphate. Biochem Biophys Res Commun 102:985–991.

10. Nissler K, Otto A, Schellenberger W, Hofmann E. 1983. Similarity of activation of yeast phosphofructokinase by AMP and fructose-2,6-bisphosphate. Biochem Biophys Res Commun 111:294–300.

11. Rodicio R, Strauss A, Heinisch JJ. 2000. Single point mutations in either gene encoding the subunits of the heterooctameric yeast phosphofructokinase abolish allosteric inhibition by ATP. J Biol Chem 275:40952–40960.

12. Gradmann D, Hansen UP, Long WS, Slayman CL, Warncke J. 1978. Current-voltage relationships for the plasma membrane and its principal electrogenic pump in Neurospora crassa: I. Steady-state conditions. J Membr Biol 39:333–367.

13. Serrano R. 1983. In vivo glucose activation of the yeast plasma membrane ATPase. FEBS Lett 156:11–14.

14. Mason AB, Allen KE, Slayman CW. 2014. C-terminal truncations of the Saccharomyces cerevisiae PMA1 H+-ATPase have major impacts on protein conformation, trafficking, quality control, and function. Eukaryotic Cell 13:43–52.

15. Rao R, Drummond-Barbosa D, Slayman CW. 1993. Transcriptional regulation by glucose of the yeast PMA1 gene encoding the plasma membrane H(+)-ATPase. Yeast 9:1075–1084.

16. García-Arranz M, Maldonado AM, Mazón MJ, Portillo F. 1994. Transcriptional control of yeast plasma membrane H(+)-ATPase by glucose. Cloning and characterization of a new gene involved in this regulation. J Biol Chem 269:18076–18082.

17. Kang WK, Kim YH, Kang HA, Kwon K-S, Kim J-Y. 2015. Sir2 phosphorylation through cAMP-PKA and CK2 signaling inhibits the lifespan extension activity of Sir2 in yeast. Elife 4.

18. Eraso P, Mazón MJ, Portillo F. 2006. Yeast protein kinase Ptk2 localizes at the plasma membrane and phosphorylates in vitro the C-terminal peptide of the H+-ATPase. Biochimica et Biophysica Acta (BBA) - Biomembranes 1758:164–170.

19. Portillo F, Eraso P, Serrano R. 1991. Analysis of the regulatory domain of yeast plasma membrane H+-ATPase by directed mutagenesis and intragenic suppression. FEBS Lett 287:71–74.

20. Lecchi S, Nelson CJ, Allen KE, Swaney DL, Thompson KL, Coon JJ, Sussman MR, Slayman CW. 2007. Tandem phosphorylation of Ser-911 and Thr-912 at the C terminus of yeast plasma membrane H+-ATPase leads to glucose-dependent activation. J Biol Chem 282:35471–35481.

21. Eraso P, Portillo F. 1994. Molecular mechanism of regulation of yeast plasma membrane H(+)-ATPase by glucose. Interaction between domains and identification of new regulatory sites. J Biol Chem 269:10393–10399.

22. Goossens A, La Fuente de N, Forment J, Serrano R, Portillo F. 2000. Regulation of yeast H(+)-ATPase by protein kinases belonging to a family dedicated to activation of plasma membrane transporters. Mol Cell Biol 20:7654–7661.

23. Rolland F, Winderickx J, Thevelein JM. 2002. Glucose-sensing and-signalling mechanisms in yeast. FEMS Yeast Res 2:183–201.

24. Ozcan S, Dover J, Rosenwald AG, Wolfl S, Johnston M. 1996. Two glucose transporters in Saccharomyces cerevisiae are glucose. Proceedings of the National Academy of Sciences 93:12428–12432.

25. Xue Y, Batlle M, Hirsch JP. 1998. GPR1 encodes a putative G protein-coupled receptor that associates with the Gpa2p Galpha subunit and functions in a Ras-independent pathway. EMBO J 17:1996–2007.

26. Viegas CA, Sá-Correia I. 1991. Activation of plasma membrane ATPase of Saccharomyces cerevisiae by octanoic acid. J Gen Microbiol 137:645–651.

27. Pontes MH, Sevostyanova A, Groisman EA. 2015. When Too Much ATP Is Bad for Protein Synthesis. J Mol Biol 427:2586–2594.

28. Browne SE. 2013. When too much ATP is a bad thing: a pivotal role for P2X7 receptors in motor neuron degeneration. J Neurochem 126:301–304.

29. Larsson C, Nilsson A, Blomberg A, Gustafsson L. 1997. Glycolytic Flux Is Conditionally Correlated with ATP Concentration in. J Bacteriol 179:7243–7250.

30. Mazón MJ, Eraso P, Portillo F. 2015. Specific phospho-antibodies reveal two phosphorylation sites in yeast Pma1 in response to glucose. FEMS Yeast Res.

31. Thomsson E, Larsson C, Albers E, Nilsson A, Franzén CJ, Gustafsson L. 2003. Carbon starvation can induce energy deprivation and loss of fermentative capacity in Saccharomyces cerevisiae. Appl Environ Microbiol 69:3251–3257.

32. Dechant R, Peter M. 2014. Cytosolic pH: A conserved regulator of cell growth? Mol Cell Oncol 1:e969643–4.

33. Thevelein JM, Hohmann S. 1995. Trehalose synthase: guard to the gate of glycolysis in yeast? Trends Biochem Sci 20:3–10.

34. Choi K-M, Kwon Y-Y, Lee C-K. 2015. Disruption of Snf3/Rgt2 glucose sensors decreases lifespan and caloric restriction effectiveness through Mth1/Std1 by adjusting mitochondrial efficiency in yeast. FEBS Lett 589:349–357.

35. Moriya H, Johnston M. 2004. Glucose sensing and signaling in Saccharomyces cerevisiae through the Rgt2 glucose sensor and casein kinase I. Proceedings of the National Academy of Sciences 101:1572–1577.

36. Flick KM, Spielewoy N, Kalashnikova TI, Guaderrama M, Zhu Q, Chang H-C, Wittenberg C. 2003. Grr1-dependent inactivation of Mth1 mediates glucose-induced dissociation of Rgt1 from HXT gene promoters. Molecular Biology of the Cell 14:3230–3241.

37. Lakshmanan J, Mosley AL, Ozcan S. 2003. Repression of transcription by Rgt1 in the absence of glucose requires Std1 and Mth1. Curr Genet 44:19–25.

38. Jouandot D, Roy A, Kim J-H. 2011. Functional dissection of the glucose signaling pathways that regulate the yeast glucose transporter gene (HXT) repressor Rgt1. J Cell Biochem 112:3268–3275.

39. Ryan OW, Skerker JM, Maurer MJ, Li X, Tsai JC, Poddar S, Lee ME, DeLoache W, Dueber JE, Arkin AP, Cate JHD. 2014. Selection of chromosomal DNA libraries using a multiplex CRISPR system. Elife 3.

40. Benjamin DI, Louie SM, Mulvihill MM, Kohnz RA, Li DS, Chan LG, Sorrentino A, Bandyopadhyay S, Cozzo A, Ohiri A, Goga A, Ng S-W, Nomura DK. 2014. Inositol phosphate recycling regulates glycolytic and lipid metabolism that drives cancer aggressiveness. ACS Chem Biol 9:1340–1350.

41. Luo B, Groenke K, Takors R, Wandrey C, Oldiges M. 2007. Simultaneous determination of multiple intracellular metabolites in glycolysis, pentose phosphate pathway and tricarboxylic acid cycle by liquid chromatography-mass spectrometry. J Chromatogr A 1147:153–164.

42. Kaiser CA, Chen EJ, Losko S. 2002. Subcellular fractionation of secretory organelles. Meth Enzymol 351:325–338.

43. Li X, Yu VY, Lin Y, Chomvong K, Estrela R, Park A, Liang JM, Znameroski EA, Feehan J, Kim SR, Jin Y-S, Glass NL, Cate JHD. 2015. Expanding xylose metabolism in yeast for plant cell wall conversion to biofuels. Elife 4.

